# Quantifying catch inequality in recreational fisheries: a case study with California steelhead (*Oncorhynchus mykiss*)

**DOI:** 10.64898/2026.03.17.712454

**Authors:** Sophia R. Sanchez, Charlie Schneider, Nann A. Fangue, Robert A. Lusardi, Andrew L. Rypel

## Abstract

Catch inequality —the disproportionate distribution of catch across anglers— is a fundamental but overlooked driver of recreational fisheries dynamics. Here, we use 11 years (2012-2022) of compulsory angler report cards to characterize long-term catch dynamics in the specialized recreational steelhead (*Oncorhynchus mykiss*) fishery in California, U.S.A. Spatialized catch data reveal the fishery is principally supported by wild fish, despite evidence of widespread hatchery straying. California steelhead appear to represent the most catch-unequal recreational fishery studied yet, exhibiting a statewide Gini coefficient of 0.81. Across basins, inequality varies (range 0.74-0.86) but remains relatively stable over time and flow conditions; high inequality is primarily driven by significant proportions of zero-catch anglers. We find the relationship between sample size and inequality measures especially influential for fisheries data. Hence, we develop a three-prong approach for identifying minimal sample sizes required for robust Gini estimation. Across basins and years, an average minimum of 77 report cards were required for the present fishery. High catch inequality may complicate traditional management efficacy and catch-per-unit-effort data utilization, emphasizing its consideration in fisheries management.

## Introduction

The tendency for catch to be disproportionately dominated by few fishers — a phenomenon known as “catch inequality” — is an innate yet underappreciated feature of all fisheries (Smith 1990; Baccante 1995). In recreational fisheries, catchability is driven by complex idiosyncratic interactions occurring between fishes and anglers (Young and Hayes 2004; Ward et al. 2013(b); Yoshiyama et al. 2023; Griffin et al. 2025). Factors influencing fish behavior and subsequent catchability (e.g. water conditions (Heermann et al. 2013; Lennox 2016), food availability (Lux and Smith 1960), and phenology) are well-explored. Angler dynamics are often highly heterogeneous across time and space as well (Hilborn 1985; Karpoff 1987; McCay and Jentoft 1996; Ward and Post 2013) yet overlooked in most studies (Salas and Gaertner 2004). Consequently, exploring catch inequality has widespread implications for understanding catchability dynamics and guiding fisheries management decisions (Cook et al. 2001; Seekell et al. 2011; Seekell et al. 2013).

Catch inequality is often quantified using the Gini coefficient, a normalized index ranging from 0-1 that describes the distribution of a resource across participants in a shared resource pool —typically wealth, given the coefficient’s economic origins (De Maio 2007). Distribution of fish catch may be quantified across various units: vessels, trips, and here, individual anglers. Though limited, current fisheries studies suggest catch inequality is rife. For example, Churchill and Snow (1964) found only 10% of anglers to be responsible for 50% of fish caught in northern Wisconsin; other contemporary fisheries express parallel patterns (Van Poorten and Post 2005; Michaletz et al. 2008; Dedual and Rohan 2016).

Catch inequality can significantly influence interpretation and utilization of common fisheries metrics, a motivating factor for managerial consideration. Catch inequality can decouple the critical assumption of a linear (ideally 1:1) relationship between CPUE and fish abundance (Post 2013). Termed “hyperstability”, this pattern manifests when CPUE fails to decline with fish populations, often due to highly skilled anglers (Hilborn and Walters 1992; Ward et al. 2013(a); Feiner et al. 2020). The rise of hyperstability in a fishery creates a high probability that CPUE measures, especially those from fishery-dependent surveys like creel surveys and report cards, are fundamentally deceptive (Dedual and Rohan 2016).

Additionally, catch inequality can unknowingly, but substantially, undermine traditional management practices. In systems with high catch inequality, strategies such as bag limits may affect only the small subset of highly skilled anglers who actually reach catch limits (Hilborn 1985; Cook et al. 2001). Further, when faced with a reduced bag limit, these anglers may simply fish more frequently, resulting in no effect of the regulation on total harvest (Camp et al. 2023). Given that bag limits are often interpreted as attainable targets, anglers catching less than set limits may perceive the fishery as unsatisfactory (Cook et al. 2001) and cease participation (Dedual and Rohan 2016); depending on catch distributions, this could be a significant proportion of participants.

Steelhead are the anadromous form of *Oncorhynchus mykiss*, and throughout California, support a socioecologically valuable fishery (Moyle 2002). There are currently six recognized Distinct Population Segments (DPS) of steelhead in the region, with five listed under the state and federal Endangered Species Acts since 1997, 1998, and 2000 (Bajjaliya 2016). Population estimates of California steelhead are rare, but most DPSs are in decline and vulnerable to extinction in the next 50 years under present conditions (Moyle et al. 2017). Analogous trends are reflected in steelhead populations across the Pacific Coast, including in Oregon, Washington, Idaho, and British Columbia (Busby et al. 2000; Kendall et al. 2017). Nonetheless, the highly popular recreational fishery for steelhead persists and is managed primarily using a long-term, compulsory angler report card system. The 31-year period of record presents an opportunity to assess long-term catch and angler dynamics across the state. This includes evaluating the role of hatchery supplementation, as hatchery-origin individuals are the only harvestable fish available for anglers since 1998 (Jackson 2007). In addition to quantifying catch inequality, the abundance of these data supports novel exploration into statistical considerations required to accurately estimate the Gini coefficient under a fisheries-data context— a necessary step in expanded use.

California steelhead are a prime example of an extremely specialized fishery with the potential for high catch inequality. Steelhead fishing is colloquially considered a highly desirable yet elusive challenge, even with existing hatchery supplementation. Spatiotemporal dynamics are key to steelhead life history and likely, their catchability and inequality (see Lux and Smith 1960; Heerman et al. 2013). Their tendency to aggregate during spawning runs (Moyle 2002) in conjunction with skilled anglers’ knowledge of when and where aggregations occur raises the specter of intense hyperstability (Erisman et al. 2011; Dedual and Rohan 2016). Therefore, we expect catch inequality to play a major role in the dynamics of this fishery, even though inequality has never been previously quantified. The goals of this study are to: (1) Characterize spatiotemporal trends in the catch of California steelhead; (2) Develop methods for estimating minimum sample sizes (angler report cards) required for reliable Gini coefficient calculation and trend assessment; (3) Quantify long-term trends in steelhead catch inequality; and (4) Explore potential seasonal and interannual drivers of catch inequality via stream flow in the fishery, a novel investigation in catch inequality literature.

## Materials and Methods

The Steelhead Report and Restoration Card (SRRC) program is an ongoing, compulsory angler report card system managed by the California Department of Fish and Wildlife (CDFW). The SRRC is designed to gather angler information pertinent to steelhead management and provide funding for enactment of such decisions (Bajjaliya 2016). Implemented in 1993 and required since 2002, the SRRC is a supplemental purchase to the state’s annual fishing license for anglers targeting steelhead (Bajjaliya 2016). Ranging from $3.15 in its conception to $9.98 in 2025, the program has sold 1.55 million report cards and generated over $7.5 million as of 2021 (Jackson 1997; CDFW 2000; Jackson 2007; Bajjaliya 2016; CDFW 2021; CDFW 2023).

From the purchase date to the annual return deadline (January 31), the SRRC requires anglers document the following information for every fishing trip targeting steelhead, regardless of catch success: Month, Day, Location Code, Wild Released, Hatchery Kept, Hatchery Released, and Hours Fished (Figure S1). Fishing on a new date or switching locations within a day constitutes a new entry (therefore after, angler trip). To minimize misidentification, the SRRC defines steelhead as “any rainbow trout greater than 16 inches found in anadromous waters” (Bajjaliya 2016). For the angler who purchased a card but did not fish for the species, there exists a “Did not fish for steelhead” checkbox. Despite the SRRC being mandatory, response rates vary annually (see below analysis testing for this effect).

Given its longevity, fishery regulations and the SRRC itself have undergone various amendments since establishment. CDFW’s 1997 initiative to adipose fin-clip 100% of hatchery individuals established basis for a statewide harvest ban of wild (non-clipped) individuals beginning in 1998 (Jackson 2007). This differentiation allowed the SRRC to classify catch by origin in 1999, formerly a single total (Jackson 2007). Reporting methods similarly changed across years: reminders via mail, telephone, and email were periodically implemented until standardization of reporting to mail and online forms in 2009 (Jackson 2007; Bajjaliya 2016). Originally 73 location codes, the SRRC eventually simplified California waterways into 20 distinct basins as of 2016 (Bajjaliya 2016) (Figure 1). We conducted our analysis on a subset of the SRRC, 2012-2022, which generally represents an extended period of methodological consistency (CDFW 2024).

**Figure 1.**
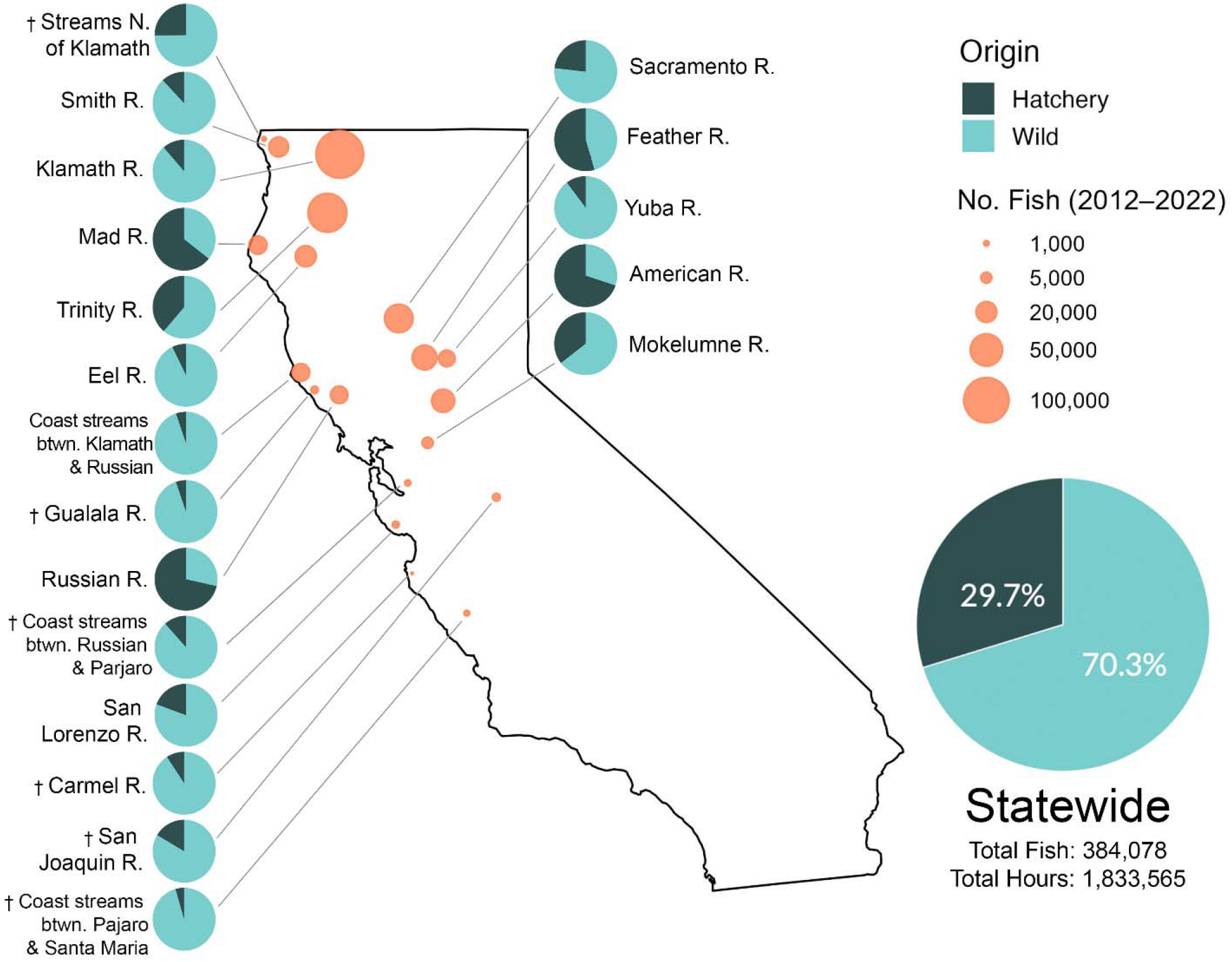
Map describing steelhead catch, 2012-2022, as reported by anglers via the CDFW SRRC Program. Inset pie charts denote the ratio of hatchery (dark blue) versus wild (light blue) origin fish for each study basin. Bubble point area proportional to sample size of steelhead caught across study period. Statewide data represents totals across all basins on the map. Dagger (†) denotes basin excluded from catch inequality analysis. California base map displayed as unprojected geographic coordinates (Latitude-Longitude, unspecified datum) from R packages “maps” (v3.4.3, Becker et al. 2025) and “mapdata” (v2.3.1, Becker et al. 2022).

To ensure anonymity, the dataset provided by CDFW scrambled unique personal angler codes (“GO ID”) but maintained the same scrambled ID per angler across angler trips within a given year; scrambled IDs were not consistent across years. Report cards featuring historical location codes were translated into the respective updated codes. SRRC angler trips with incomplete fish count or invalid location codes were excluded from the dataset (5.72% of angler trips). The resulting dataset featured 431,450 angler trips catching a total of 384,078 individual steelhead (hatchery and wild, kept and released), as reported by an annual average of 5,629 – 9,242 anglers (2022 and 2012, respectively). Calculations of CPUE further excluded angler trips with unreported “Hours Fished” (3.26% angler trips). Characterization of catch trends, including number of fish caught, CPUE, and wild-to-hatchery ratios, was conducted on a basin-specific level using data aggregated from 2012-2022. All location codes (hereafter referred to as “basins”) were included in this analysis with the exception of the southernmost basin (“Santa Maria River to the Mexican border”) due to its closure to steelhead fishing during the study period. Catch trend maps were created using the “mapdata” (v2.3.1, Becker et al. 2022) and “maps” (v3.4.3, Becker et al. 2025) R packages. All following statistical analyses were conducted using R Statistical Software v4.4.1 (R Core Team 2025) with most visuals generated using the “ggplot” R package (v4.0.0, Wickham 2016).

We characterized catch inequality on an annual, basin-specific basis using Lorenz curves and Gini coefficients (Smith 1990; Baccante 1995; Cook et al. 2001; Van Poorten and Post 2005; Michaletz et al. 2008; Seekell et al. 2011; Seekell et al. 2013; Dedual and Rohan 2016). Lorenz curves plot the cumulative percent of anglers by the cumulative number of fish caught, in comparison to a 1:1 line representing perfect equality (Smith 1990; Baccante 1995) (Figure 2). Deviations from perfect equality are quantified by the Gini coefficient— traditionally used in economics and best known for its calculations of global income inequality (De Maio 2007). Gini is computed as the area between the 1:1 line and the Lorenz curve (*H*), divided by the total area under the 1:1 line (*H / H+h*) (Smith 1990; Baccante 1995). A Gini of 0 indicates maximal equality in catch (as the Lorenz curve is 1:1) whereas a Gini of 1 demonstrates maximal deviation from equality (Smith 1990; Baccante 1995). These metrics were repeated for calculations of harvest inequality, which only considers the subset of fish caught and thereafter harvested (e.g. not released). For all Gini calculations and subsequent analysis, we summarized angler trips such that individual anglers were represented once per basin annually (e.g. annual sum of an angler’s total catch and effort per basin). This unit (henceforth, “angler report card” for a given basin and year) allows for spatiotemporal analysis of catch inequality at the individual angler level by (1) preventing replicate representation of anglers who report multiple trips to the same basin within a given year, while (2) treating each year as a unique composition of anglers, which may contain both returning and new participants. Following this, all angler trips condensed into 121,505 angler cards. Calculations of Gini and Lorenz curves were performed using the “DescTools” R package (v0.99.6, Signorell 2025). Differences in annual Ginis across basins and regions were assessed using a Kruskal-Wallis test (“rcompanion” R package v2.5.0, Salvatore 2025) followed by the post-hoc Dunn’s test (“FSA” R package v0.10.0, Ogle et al. 2025) to account for the non-parametric nature of Gini (Zhang et al. 2020).

**Figure 2.**
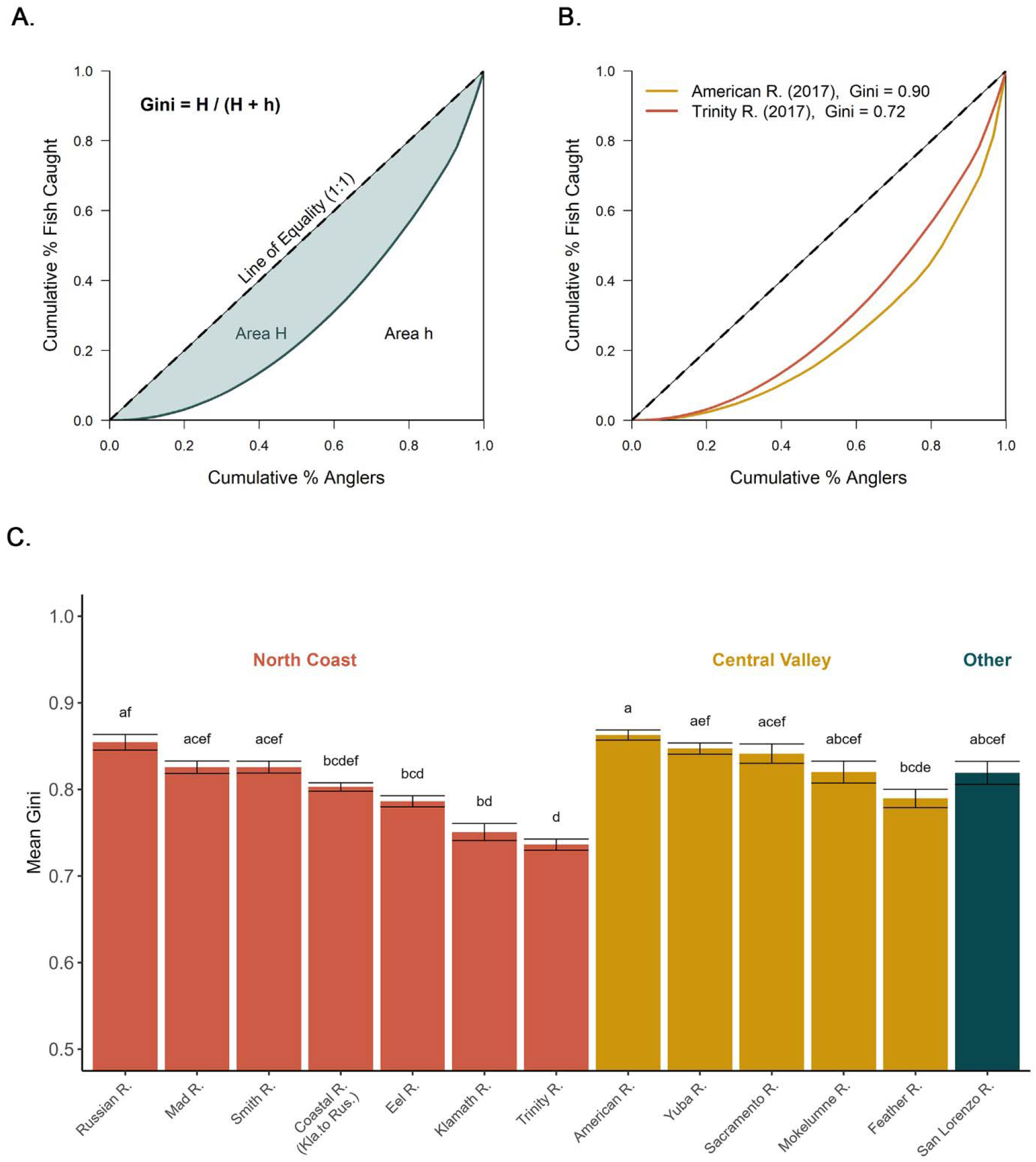
Method and results of Gini coefficient calculations. Basin bar coloration signifies geographic region: North Coast (red), Central Valley (yellow), or Other (deep blue). (A)Demonstration of Gini estimation using Lorenz curves. The closer the Lorenz curve (blue line; determined by resource distribution) is to the Line of Equality (dashed line; maximal equality in resource distribution), the lower the inequality (Gini) of a system. (B) Example calculations of Gini values for two study basins in the same year. Inequality is higher in the American River (yellow). (C) Mean Gini coefficients by basin across all years (2012-2022) +/- 1 SE. Basins sharing same letter above standard error bars are not statistically distinct from one another (Kruskal-wallis and Dunn’s post hoc test, P > 0.05).

While the Gini coefficient offers a numerical representation of catch inequality, it does not provide context on its drivers. As such, we calculated the Lorenz asymmetry coefficient (LAC) to identify the relative contributions of high-catch vs low-catch anglers to Gini patterns (“ineq” R package v0.2-13, Zeileis 2014). A LAC of 1 indicates equal contributions to the Gini, whereas LAC >1 demonstrates skew driven by the high catch of few, skilled anglers. Conversely, where LAC is < 1, skew is driven by numerous low-to-zero catch anglers (Dedual and Rohan 2016). Correlations between LAC and Gini were tested (Seekell et al. 2013; Dedual and Rohan 2016) using Spearman’s correlation coefficient and Bonferroni corrections (Zhang et al. 2020).

Although relatively unexplored under a fisheries context, economic literature of the Gini coefficient often mentions sample size biases (Deltas 2003). Given the large variations in sample size (here, number of angler report cards) across basin-year combinations, we developed a three-pronged approach to ensure Gini calculations and trends were not influenced by sample size (Deltas 2003). To determine the minimum sample size required for reliable Gini computation, a power analysis comparing sample size to Gini was performed. For every basin-year combination, angler report cards were randomly subset into groups of *n* = 25 and Gini calculated for every group (Dixon et al. 1987). Using a broken-stick model (“segmented” R package v2.1-4, (Muggeo 2008)), a “breakpoint” signifying the minimum sample size required for estimation of a stable Gini was calculated. This process was repeated recursively such that 500 minimum breakpoint estimates were calculated (Dixon et al. 1987). Using the 1st quartile value of these 500 breakpoint estimations, we computed a final statewide breakpoint and basin-specific breakpoints to account for variation in sample size across basins and within years. The 1^st^ quartile provided a reasonable but lenient estimate of minimum sample size. This was done, in part, to better contextualize our results with those found in the literature; some of which have quite low samples (e.g., N < 3-5). If we were to use the median or 3^rd^ quartile estimates, the minimum sample size would increase considerably more, and some moderately low sample basins in our study would likely be removed. Basins with significantly low samples sizes failed the initial breakpoint analysis altogether, never reaching a stable estimation within the iterations. Therefore, our thresholds should truly be considered the truly minimum possible sample bar, at least in our dataset, but likely others as well. To be considered for analysis, every year in a basin must have satisfied both the statewide and basin-specific breakpoints (e.g. exceeded the minimum number of angler cards required across all basins, and specific to the respective basin). Following identification of sites with sufficient sample size for reliable Gini, we conducted a general linear model (GLM) to test whether response rate influences Gini at a statewide scale. These models cannot be executed at a lower spatial resolution because response rates are only available at an annual, statewide scale. In this GLM, mean annual statewide Gini (calculated as average across-basin Gini for each year) was the dependent variable, and response rate was the independent variable. Normality tests for annual statewide and basin-specific Ginis were performed prior to executing the model.

Finally, we conducted a linear mixed-effects regression (R packages: “lme4” v1.1-37 (Bates et al. 2015); “lmerTest” v3.1-3 (Kuznetsova et al. 2017); “MuMIn” v1.48.11 (Bartoń 2025)) to assess relationships between sample size and Gini, and subsequently used this model to detrend each Gini estimate from the influence of sample size. In the model, Gini was the response variable, log_10_(sample size) was the fixed effect, and log_10_(sample size) and basin were random effects. As such, log_10_(sample size) was used as a fixed and random effect (*Gini ∼ log(number angler report cards) + (log (number angler report cards)| basin))*).Our primary goal was to “detrend” Gini from interannual variations of n with each basin (e.g., random effects); thus, we did not extrapolate results of the parent regression for prediction. Residuals from this model were plotted over time to assess directional trends in Gini across basins.

We used variation in stream flow to assess potential seasonal drivers of catch inequality. Flow data were downloaded from U.S. Geological Survey (USGS) stream gages and assessed for bivariate correlation with Gini and LAC using Spearman correlations. To address the potential of multiple comparisons, we adjusted the standard alpha of 0.05 using the Bonferroni correction (Zhang et al. 2020). For each basin, we used the watershed’s lowermost USGS gauge with sufficient time coverage (Table S1). Based on these criteria, flow data were available for the following rivers: Smith, Klamath, Trinity, Mad, Eel, Russian, San Lorenzo, Sacramento, Yuba, American, and “Other coastal streams and rivers between the Klamath and Russian Rivers” (USGS 2024). We repeated all aforementioned analyses on different subsets of the data (e.g., wild-vs-hatchery catch, and seasonal catch subsets representing peak fishing months). The number of angler report cards, when grouped by fish origin, did not meet the pre-determined minimum sample size thresholds, and no seasonal differences were observed in the analysis; therefore, only annual total catch results are presented for simplicity.

To compare catch inequality for California steelhead to other fisheries, we conducted a systematic review of studies examining Gini in other recreational fisheries. Only studies with an annual sample size (aggregated across locations) greater than our minimum statewide *n* threshold were considered sufficient to avoid sample size influence on Gini and were thus retained. Other studies were rejected from these comparisons. In cases where only a multi-year, aggregated sample size was provided, it was divided by the number of study years as a proxy for annual sample size. Similarly, where both a Gini value for all anglers and only anglers targeting specific species were provided, only the targeting Gini was considered (Cook et al. 2001).

## Results

Total numbers of fish caught and origin of steelhead catch (wild or hatchery) varied widely across basins; however, CPUE did not (mean = 0.14, SD = 0.07) (Figure 1). Catch demonstrated a strong latitudinal gradient in California, with northern regions posting higher catch totals (e.g., North Coast *n* = 267,732) than central basins (e.g., Central Valley *n* = 112,046). Combined catch in the most southerly regions was lower (e.g., Southern California *n* = 4,300). The proportion of wild-to-hatchery steelhead in each basin was related to hatchery presence; basins maintaining hatchery facilities predictably had the highest percentage of hatchery-origin fish (Table 1). Notably, hatchery-origin fish were reported in all 19 study basins (ranging 4-71% of catch) despite most basins lacking any hatcheries. Within these hatchery-free basins, hatchery fish accounted for 4-25% of catch across all years. Nonetheless, on a statewide-basis, wild-origin fish comprised 70.3% of catch over the full study period.

**Table 1.** Summary information on steelhead data captured through the California SRRC. Data reflect all years, 2012-2022. Mean % hatchery represents average annual percentage of hatchery-origin catch per basin; total range provided in Range % hatchery. Minimum straying estimate only accounts for hatchery-to-non-hatchery straying and therefore cannot be calculated for hatchery basins, as indicated by n.a.

|  | **Basin** | **Total fish** | **Range % hatchery** | **Mean % hatchery** | **Mean minimum stray estimate (%)** |
| --- | --- | --- | --- | --- | --- |
| Non-Hatchery | Smith R.<sup>a</sup> | 18058 | 7.7 - 16.4 | 11.9 | 11.9 |
|  | Coastal R. 1<sup>b</sup> | 630 | 6.0 - 60.0 | 25.1 | 25.1 |
|  | Eel R. | 14849 | 2.9 - 12.7 | 7.2 | 7.2 |
|  | Coastal R. 2<sup>c</sup> | 14108 | 1.5 - 9.55 | 5.2 | 5.2 |
|  | Gualala R. | 2024 | 0.0 - 33.3 | 5.2 | 5.2 |
|  | Yuba R. | 12161 | 6.0 - 19.6 | 10.3 | 10.3 |
|  | Coastal R. 3<sup>d</sup> | 1326 | 2.7 - 21.4 | 11.5 | 11.5 |
|  | San Joaquin R. | 2346 | 4.9 - 37.2 | 16.4 | 16.4 |
|  | San Lorenzo R.<sup>a</sup> | 1779 | 3.3 - 34.8 | 19.3 | 19.3 |
|  | Carmel R. | 87 | 0.0 - 31.6 | 9.2 | 9.2 |
|  | Coastal R. 4<sup>e</sup> | 1103 | 0.0 - 11.0 | 4.4 | 4.4 |
| Hatchery | Klamath R.<sup>f h</sup> | 110664 | 7.05 - 20.0 | 11.3 | n.a. |
|  | Trinity R.<sup>f</sup> | 73148 | 27.5 - 47.5 | 38.8 | n.a. |
|  | Mad R.<sup>f</sup> | 20454 | 52.3 - 73.4 | 64.4 | n.a. |
|  | Russian R.<sup>f</sup> | 13797 | 61.2 - 82.7 | 71.5 | n.a. |
|  | Sacramento R. <sup>g</sup> | 38610 | 14.4 - 40.7 | 23.2 | n.a. |
|  | Feather R.<sup>f</sup> | 28745 | 34.4 - 66.2 | 54.5 | n.a. |
|  | American R.<sup>f</sup> | 24937 | 60.0 - 82.0 | 69.9 | n.a. |
|  | Mokelumne R.<sup>f</sup> | 5247 | 21.8 - 52.7 | 35.6 | n.a. |

<sup>a</sup> Private hatchery present, but considered non-hatchery due to unclear marking practices.
<sup>b</sup> Coastal St./R. north of Klamath.
<sup>c</sup> Coastal R. between Kla. R & Rus. R.
<sup>d</sup> Coastal St./R. between Russian and Pajaro R.
<sup>e</sup> Coastal St./R. from Pajaro (incl.) to Santa Maria R.
<sup>f</sup> CDFW hatchery.
<sup>g</sup> USGS hatchery.
<sup>h</sup> Iron Gate Hatchery included, but closed in 2021.

Completion of the breakpoint and power analysis revealed the statewide minimum sample size breakpoint (mean number of all annual angler report cards) required for stable Gini calculations to be 77. When assessed on a basin-specific scale, breakpoints ranged 60-141 (Table 2). While annual return rate varied (range 26-43%, mean = 34.6%), there was no significant relationship between annual statewide Gini and annual return rate (GLM, df = 8, F = 0.05, *p* = 0.82, R^2^ = 0.007). Application of basin-specific minimum sample size breakpoint to our full dataset resulted in disqualification of 7 of 19 basins (37%) for further analysis. Specifically, we eliminated “Coastal rivers north of the Klamath”, the Gualala, “Coastal rivers between Russian and Pajaro rivers”, the Carmel, the San Joaquin, “Coastal rivers between Pajaro and Santa Maria rivers” basins because of insufficient sample size (Table 2). The 13 basins with suitable numbers of angler report cards were the Russian, Mad, Smith, Eel, Klamath, Trinity, American, Yuba, Sacramento, Mokelumne, Feather, and San Lorenzo Rivers, as well as the “Coastal streams between Klamath and Russian Rivers” basin. The final dataset included data from a total of 115,926 angler cards collected during 2012-2022.

**Table 2.** Minimum *n* thresholds by basin and statewide average, as determined by Gini power analysis. The card count range indicates annual sample size, or number of returned SRRC entries across 2012-2022. Minimum *n* thresholds represent the first quartile summary of all annualized thresholds by basin. n.a. indicates no breakpoint during simulations due to low sample size.

| **Basin** | **Minimum $n$ threshold** | **Card count range** |
| --- | --- | --- |
| *Statewide* | 77 | 13 - 2625 |
| Trinity R. | 141 | 1444 - 2625 |
| Klamath R. | 130 | 1170 - 1906 |
| Sacramento R. | 104 | 695 - 1633 |
| American R. | 101 | 718 - 1433 |
| Coastal R. 4<sup>a</sup> | 84 | 48 - 169 |
| Russian R. | 79 | 424 - 1152 |
| Mad R. | 78 | 440 - 870 |
| Eel R. | 78 | 418 - 936 |
| Coastal R. 2<sup>b</sup> | 78 | 229 - 650 |
| Smith R. | 77 | 334 - 1059 |
| Yuba R. | 77 | 205 - 537 |
| Feather R. | 76 | 471 - 815 |
| Gualala R. | 70 | 46 - 179 |
| San Lorenzo R. | 68 | 83 - 204 |
| San Joaquin R. | 68 | 64 - 146 |
| Mokelumne R. | 67 | 95 - 242 |
| Coastal R. 3<sup>c</sup> | 60 | 60 - 152 |
| Coastal R. 1<sup>d</sup> | n.a. | 21 - 86 |
| Carmel R. | n.a. | 13 - 50 |

<sup>a</sup> Coastal St./R. from Pajaro (included) to Santa Maria R.
<sup>b</sup> Coastal R. between Kla. R & Rus. R.
<sup>c</sup> Coastal St./R. between Russian and Pajaro R.
<sup>d</sup> Coastal St./R. north of Klamath

Catch inequality across the California recreational steelhead fishery was extreme (statewide mean Gini = 0.81, SD +/- 0.05). Basins expressed differences in Gini, with values ranging 0.74-0.86 (Figure 2, Table S2). In general, catch inequality was higher in the Central Valley (mean = 0.83, SD +/- 0.04) versus the North Coast (mean = 0.80, SD +/- 0.05). This difference between regions was significant (Kruskall-Wallis and Dunn’s Post-Hoc, df = 2, chi-squared = 18.2, adjusted *p* = 0.00006). For comparison purposes, we quantified statewide harvest inequality which was predictably found to be more extreme than catch inequality (mean = 0.94, SD +/- 0.08) and varied across basins similarly (range = 0.86 - 0.99).

The global effect of sample size on Gini across basins was negative but not significant (Figure 3, Mixed Effect Model, t = -1.82, *p* = 0.12, pseudo-R^2^ marginal= 0.09); when we allowed this effect to vary by location, the magnitude of impact increased dramatically (pseudo-R^2^ conditional= 0.80). Therefore, we used the relationship specific to each basin to detrend each Gini estimate from the influence of sample size. Following detrending, there were almost no temporal relationships between detrended Gini (residuals) and year for any of the study basins (Figure 3). Catch inequality was temporally stable in the Russian, Mad, Smith, Coastal rivers between Klamath and Russian, Eel, American, Yuba, Sacramento, Mokelumne, Feather, Trinity, and San Lorenzo Basins (GLMs, all dfs = 9, all *p’s* >0.003 following Bonferroni correction). The one exception was the Klamath River Basin, which expressed decreasing catch inequality over our study period (GLM, *p* = 0.001).

**Figure 3.**
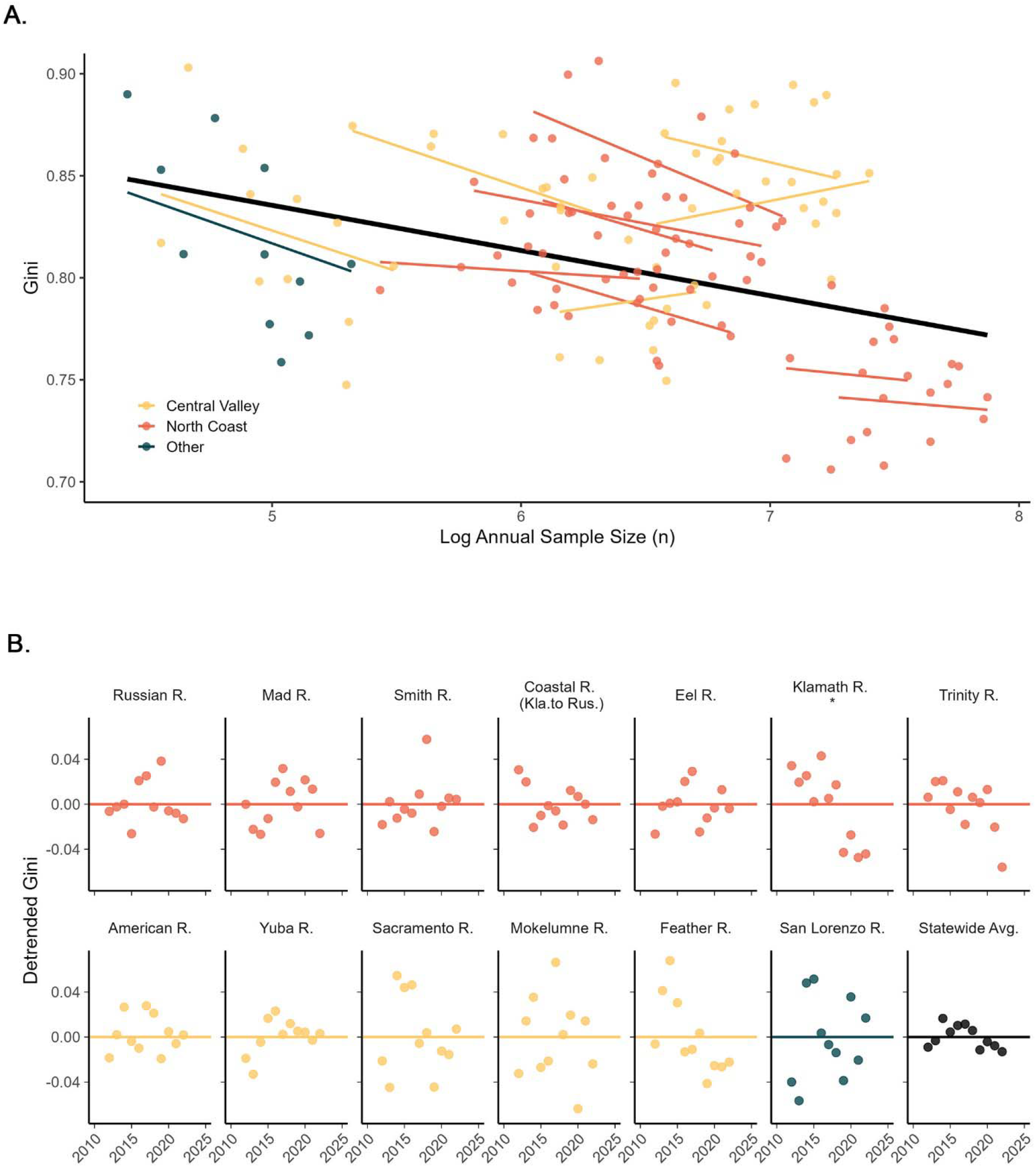
Examining the relationship between Gini, sample size, and year. Bar coloration signifies geographic region: North (red), Central Valley (yellow), Other (deep blue), and Statewide (black). (A) Relationship between annual steelhead Gini and sample size (card count) in each study basin, as represented by: *Gini ∼ log(Number Angler Report Cards) + (log(Number Angler Report Cards)| Basin))*. Thick black line denotes parent regression for the mixed effect regression model. Thin colored lines denote random effects, e.g. basins. (B) Detrended Gini plotted over time in each study basin. Detrended Gini is calculated as residuals from the mixed effects model presented in A. Horizontal line reflects a neutral value of zero. There were no significant correlations between detrended Gini and year, with the exception of the Klamath Basin (as denoted by asterisk).

LAC similarly showed slight variations across regions, basins, and time, ranging 0.79-0.93 (mean = 0.88, SD +/- 0.07) (Table S2). However, following the LAC categories proposed by Dedual and Rohan (2016), there was little difference in result interpretations, suggesting zero-catch anglers as the dominant driver of inequality across basins. Indeed, at the basin level, the average annual proportion of anglers catching zero fish demonstrated a positive relationship with catch inequality (Table S2). The least catch unequal basins, Klamath and Trinity Rivers, exhibited the lowest percentage of zero-catch anglers (37.0%, and 40.8% respectively), while more extreme catch unequal basins like the Russian River exhibited highest percentages (68.7%). Unlike the findings of previous studies, Gini and LAC were uncorrelated (multiple R^2^ = 0.0004, *p* = 0.81).

In every study basin with available flow data, Gini and LAC estimates were largely uncorrelated with annual flow (Gini Spearman Correlation for Gini and Flow, *p* range: 0.05 - 0.88, Gini Spearman Correlation for LAC and flow, *p* range = 0.08 - 0.94) (Figure 4). No values approached significance following Bonferroni adjustments. These relationships were also tested on a monthly timestep, but no significant correlations were found between monthly flows and either Gini (Spearman Correlation, *p* range = 0.04 - 0.94) or LAC (*p* range = 0.01 - 0.94), again after applying the Bonferroni-adjusted D level.

**Figure 4.**
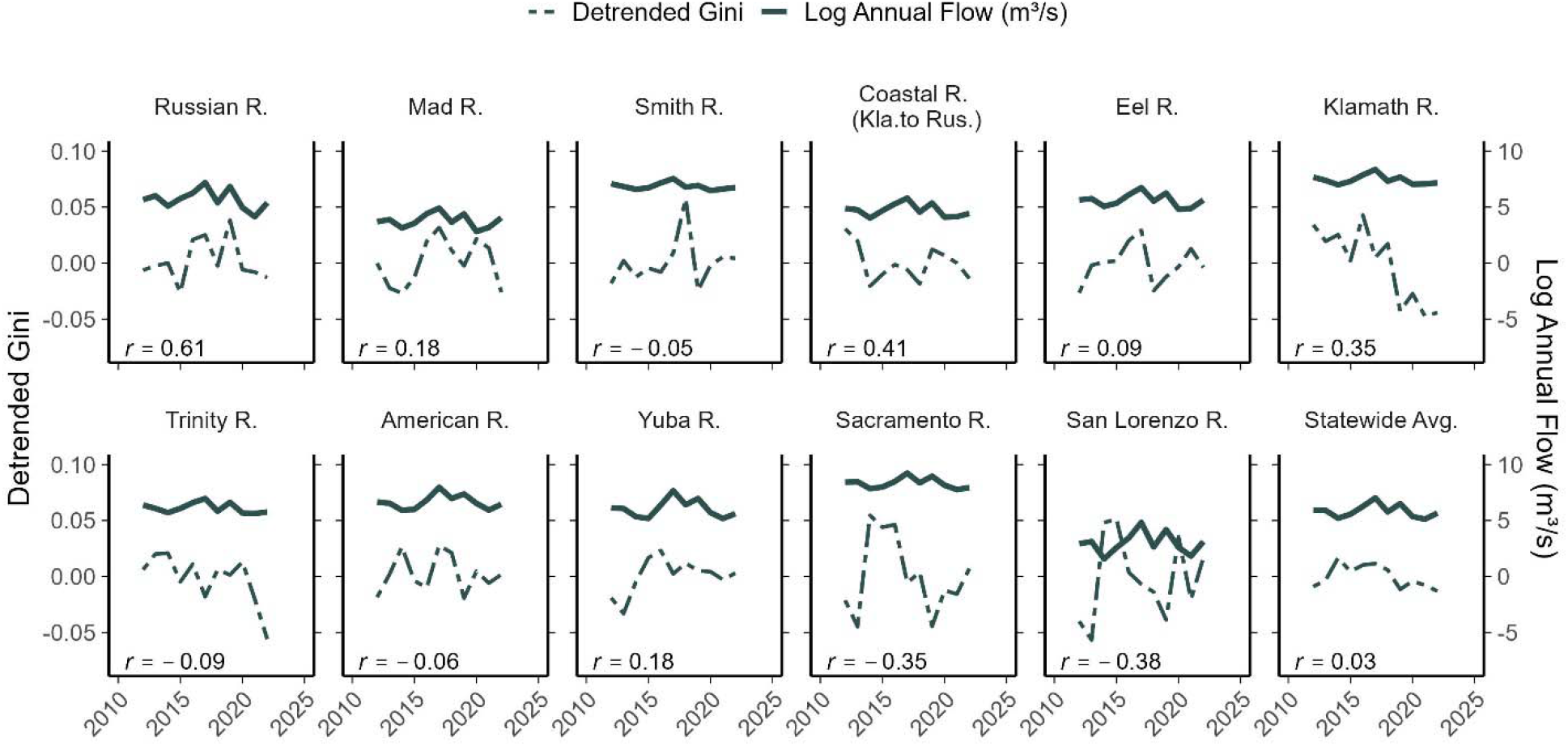
Relationship between detrended Gini (dashed lines) and log annual flow (cubic meters per second; solid lines) by basin, 2012-2022. Statewide average is calculated as the mean Gini coefficient across our study basins for each year, and statewide flow also represents only these same basins. Spearman correlations for each basin (r values) indicate no significant correlations (P > 0.05) between inequality and flow.

## Discussion

Here, we explore angler-generated data for 11 consecutive years on recreational steelhead catch trends and inequality across freshwater ecosystems in California, presenting findings as valuable considerations for their sustainable management. First-order catch characterizations revealed wild-fish dominated steelhead catch on a statewide scale (70.3% total catch) despite concurrent evidence of hatchery-fish straying across California basins, underscoring the importance of prioritizing wild steelhead production in management (Katz et al. 2013; Huber et al. 2024). Hatchery-origin steelhead were reported in every assessed basin with varying degrees, averaging 11% even in hatchery-free basins (Table 1). Hatchery catch estimates presented here indicate the widespread prevalence, and subsequent influence, of hatcheries on wild steelhead populations. These findings are consistent with prior studies (Araki and Schmid 2010; Sass et al. 2017; McMillan et al. 2023) and with research on Pacific salmonids specifically (Knudsen et al. 2021; Dayan et al. 2024; Riddell et al. 2024). Across Washington rivers, hatchery-origin steelhead comprised 15-89% of catch (Dauer et al. 2009), and in hatchery-free basins in Oregon, 22% (Schroeder et al. 2001). While the ecological effects of hatchery fish vary (Berejikian and Ford 2004; Araki et al. 2007), even progressive hatchery practices can impact wild populations — for example, by slowing growth, increasing predation, or inducing early migration in wild individuals (Kostow 2009; Hayes et al. 2013). Anthropogenic reductions in access to headwaters, wetlands, and estuaries (Lindley et al. 2006), along with altered flow regimes (Yoshiyama et al. 2001), similarly threaten California steelhead (Moyle et al. 2017). Given the recreational fishery’s reliance on wild populations, addressing these conservation concerns is likely crucial to its long-term persistence.

Geographic trends in catch support the strong, positive latitudinal gradient in steelhead abundance documented within California. In general, freshwater fishes often demonstrate marked latitudinal trends in growth (Conover and Present 1990; Pegg and Pierce 2001; Rypel 2012), body size (Belk and Houston 2002; Rypel 2014), biomass and productivity (Rypel and David 2017; Myers et al. 2018). While cold-water species often exhibit the strongest latitudinal patterns (Rypel 2012; Rypel and David 2017), these trends are still somewhat understudied, especially along the Pacific Coast (Myers et al. 2018; Cano-Barbacil et al. 2023; but see Nicola et al. 2009). Accordingly, in California, steelhead populations south of Point Conception are notably smaller than those in northern basins (Garza et al. 2014), a hypothesized consequence of southern California water temperatures approaching steelhead thermal tolerance limits (Light et al. 1989; LaDochy et al. 2007; Katz et al. 2013), and/or decreased cold-water habitat availability (Busby et al. 1996; Katz et al. 2013). Total catch across our study period followed a similar trend, with most catch occurring in most northward basins and decreasing with latitude.

When calculated at the state level, steelhead catch inequality in California, was among the most unequal recreational fisheries ever documented (Table 3). While all recreational fisheries exhibit some level of intrinsic catch inequality (Baccante 1995; Cook et al. 2001; Seekell et al. 2013), the extreme statewide Gini coefficient of 0.81 (range 0.73-0.86, by basin) indicates that a small number of highly skilled anglers account for the vast majority of the fish caught. Although comparable studies on other steelhead fisheries are lacking, research on recreational rainbow trout fisheries (*O. mykiss*, as well) also demonstrate catch inequality, though generally not as pronounced. In Alberta, the first 2 years of a newly stocked lake exhibited an average Gini coefficient of 0.67 (range 0.48-0.81, over time) (Van Poorten and Post 2005). In New Zealand’s Tongariro River basin, naturalized rainbow trout populations demonstrated an average Gini coefficient of 0.58 (Dedual and Rohan 2016). Despite stark differences in catch distribution, high proportions of low-catch anglers in both the Tongariro River basin rainbow trout fishery and California steelhead fishery primarily drove respective inequalities (Dedual and Rohan 2016).

**Table 3.**
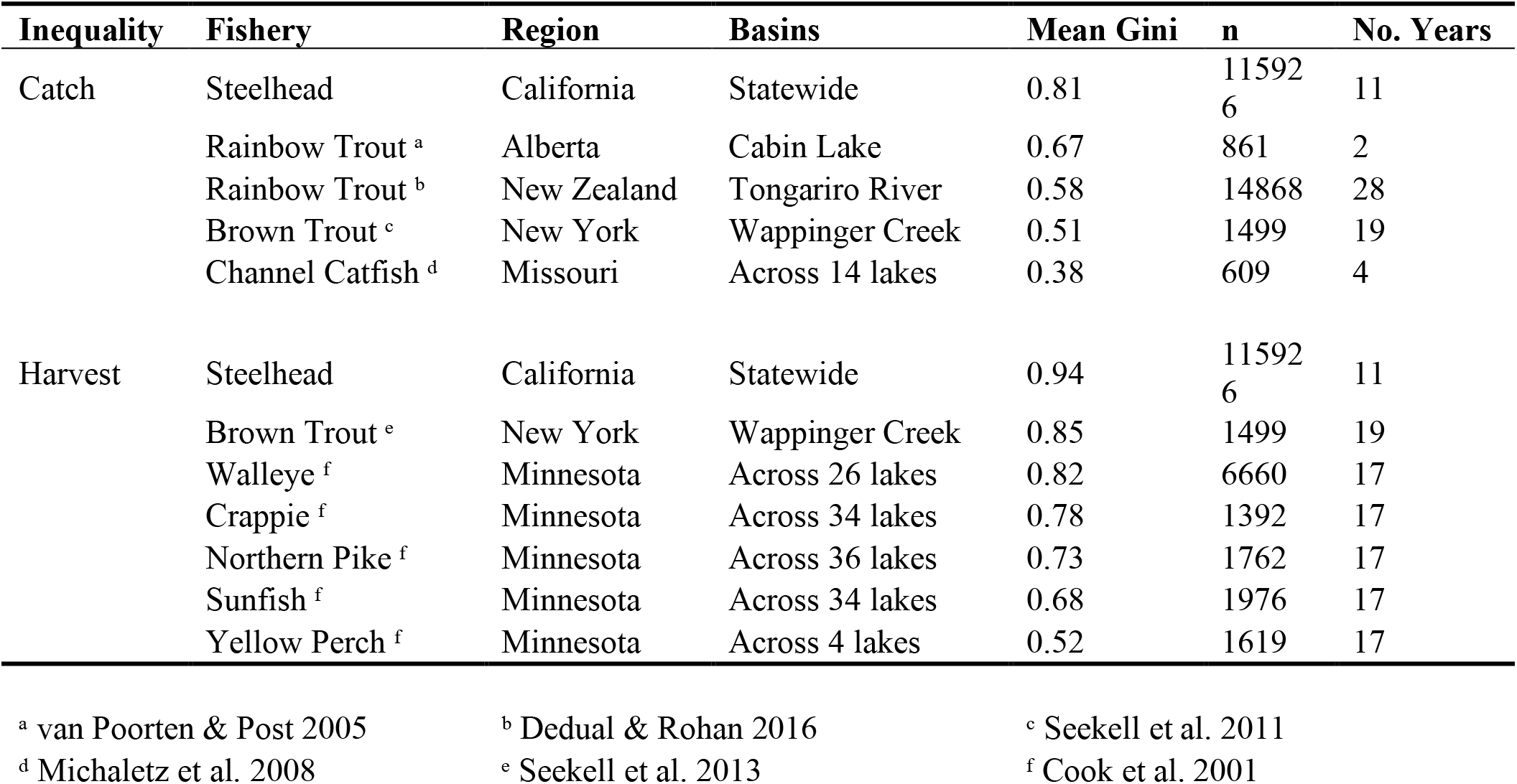
Comparison of California steelhead catch and harvest inequality estimates to other recreational fisheries from the literature.

Catch and harvest inequality proved similarly muted across other species and fisheries. A brown trout (*Salmo trutta*) fishery in New York exhibited an average Gini coefficient of 0.51 (Seekell et al. 2013), while stocked channel catfish (*Ictalurus punctatus*) in “put-grow-take” impoundments demonstrated the lowest catch inequality of measured systems with an average Gini coefficient = 0.38) (Michaletz et al. 2008). Occasionally, studies quantify harvest inequality to evaluate the effectiveness of harvest regulations (Cook et al. 2001; Seekell et al. 2013). From first principles, we expect harvest inequality to be at least as high—and likely higher—than catch inequality, because only a subset of caught fish are harvested. For instance, the New York brown trout fishery exhibited a harvest inequality of Gini = 0.85, notably higher than its corresponding catch inequality (Seekell et al. 2013). We therefore can conclude that any studies presenting harvest Gini inequalities, e.g., Walleye *Sander vitreus* (0.82), Crappie *Pomoxis nigromaculatus* (0.78), Northern Pike *Esox lucius* (0.73), Sunfish *Lepomis spp*. (0.68), and Yellow Perch *Perca flavescens* (0.52) (Cook et al. 2001) likely demonstrate higher values than their respective catch Gini coefficients. Thus, California steelhead catch inequality is apparently the highest Gini estimated to date for catch using robust sample sizes.

Even so, our steelhead Gini coefficient may still be an underestimation of the fisheries’ true catch inequality. Statewide and basin-specific LACs reveal the high percentage of specifically zero-catch anglers as the principal driver of the elevated Gini coefficient; on average, 45.8% of anglers will fail to catch a single fish within a year, regardless of basin. However, demographic studies of the SRRC program document avid anglers (e.g., anglers that are older, frequently repurchase SRRCs, hold lifetime fishing licenses) as more likely to return report cards (Gusman Costa and Hause 2023; see also Pollock et al. 1994; Dorow and Arlinghaus 2011). Given that the drivers of our already extreme Gini coefficient—lower skilled anglers—are underrepresented in our data, the true Gini coefficient of this fishery is likely even higher.

A key finding from this study is the extent to which the Gini coefficient is strongly influenced by angler report card sample size, a critical consideration when precisely quantifying catch inequality particularly in recreational fisheries. The influence of sample size on the Gini coefficient has been reported previously; however, to our knowledge it has not been comprehensively addressed in the fisheries literature until now. Existing attempts (see Seekell et al. 2011; Dedual and Rohan 2016) account for small *n’s* by utilizing unbiased Gini formulas common in the economics literature (Glasser 1962; Yao 1999). Unbiased Gini coefficient calculations attempt to correct for systematic, downward biases often observed in economic applications with small sample size (Deltas 2003) by ordering data by an increasing metric (here, effort) and multiplying results by (*sample size)/[(sample size)-1]* (Glasser 1962; Dixon et al. 1987; Yao 1999; Damgaard and Weiner 2000). However, as evidenced by our mixed-effects model, the influence of sample size on the Gini coefficient is variable. The catch Gini coefficient showed that its values can vary by location, and subsequently angler populations, with extremely high and low values often occurring when the number of angler report cards are small. This is likely because the structure of the fishery data is fundamentally different from economics data: zero-catch anglers are often comprise significant proportions of the angler population whereas in human societies most individuals earn some (non-zero) income. Consequently, fisheries data are potentially vulnerable to inflated Ginis at low sample sizes because of the increased abundance of zero-catch anglers and lower probability of encountering a highly skilled angler. Sample size issues may be explored by assessing Gini at an alternative scale (e.g. trip, vessel) to the individual angler, with respectively adjusted interpretation. Where differential data grouping is not possible or of interest, increasing sample size complexly requires recruitment of anglers into the fishery, and/or improved response rates. Indeed, angler-reported datasets often have high and variable non-response rates, an admittedly difficult challenge to overcome (Pollock et al. 1994; Gigliotti and Fopma 2019). We therefore recommend that future studies of fisheries’ catch inequality explicitly consider sample size and basin-specific angler demographics (including temporal change) when quantifying Gini.

Factors proposed to influence catch inequality can be broadly categorized as managerial (gear advancements and restrictions, regulation strategy), environmental (abiotic and biotic conditions impacting fish abundance, behavior, etc.), or chance (Van Poorten and Post 2005; Dedual and Rohan 2016). Complexly, angler behaviors may interact with any of these variables. As such, temporal catch inequality trends vary: while brown trout demonstrated long-term stability (Seekell et al. 2011), established rainbow trout fishery Gini decreased over time (Dedual and Rohan 2016), and increased in newly stocked fisheries (Van Poorten and Post 2005). Disentangling the causal mechanisms in catch inequalities across species is beyond the scope of this study. However, we explored these mechanisms across basins and years in the California steelhead fishery. After detrending the Gini coefficient for inter-annual sample size variations, we found annual catch inequality and LAC to be temporally stable, except for one basin (Klamath). Lack of notable temporal variation in both metrics may help explain why no correlation was detected between Gini and LAC, as restricted value ranges can reduce the ability to observe any underlying association. To test environmental hypotheses, all analysis were repeated on a subset of data only featuring each region’s respective fishery season, though results were not significantly different. We further explored potential relationships between flow and Gini at a variety of temporal scales but found no relationship. In some basins, steelhead fishing is temporarily prohibited under basin-specific, low flow thresholds; potential closure impacts were not analyzed as data aggregated on a daily time scale failed to meet minimum sample size requirements. Seasonality and flow therefore did not play a strong role in the Gini estimation.

We suggest steelhead catch inequality and its spatial variation is most likely a product of unmeasured dynamics of angler behavior, a topic that may be of interest for future research. A variety of factors within the steelhead fishery changed over the period of this study, such as the transition from physical to virtual dominant reporting method, and the inevitable evolution of angling gear. However, the observed stability in both overall angler skill and skill-class suggest that no one source fundamentally altered the distribution of catch across anglers. Similarly, the consistent decline in angler participation over these 11 years failed to significantly shift the driving demographic of catch inequality from low-catch anglers. Discussions with California steelhead anglers indicate that differential prevalence of guiding across basins (and more broadly, fisheries) may influence Gini as the sharing of guide knowledge and skill (Farthing et al. 2022) reduces skill differential. Indeed, basins anecdotally identified as having strong guiding presence (e.g. Klamath and Trinity) demonstrated lower catch inequality than the statewide average. However, information on guiding is not currently collected as part of the SRRC. Future incorporation of guide usage by trip on the SRRC could better assess the role guiding plays in catch dynamics, and its subsequent impacts on data analysis and management decisions by basin.

While interactions between angler dynamics and population characteristics may further illuminate steelhead catch inequality, examining these relationships underscores a more fundamental issue: the critical consideration of catch inequality interpreting angler-based CPUE as a measure of relative abundance. Where total steelhead catch per basin exhibited a positive latitudinal trend, CPUE did not, highlighting a likely influence of angler behavior and skill (Dabrowksa et al. 2017; Feiner et al. 2020; Mosley et al. 2022). Indeed, the increased knowledge, experience, and/or resources of high-catch anglers may allow for success even in lower density steelhead basins, thereby obscuring spatial patterns in CPUE. Basin-level Gini values, when compared regionally, largely validate this hypothesis. More southern Central Valley basins, where steelhead densities are lower, exhibited higher levels of catch inequality, suggesting that greater angler skill is required to achieve success. Similar increases in catch inequality under conditions of low fish abundance have been documented previously (Baccante 1995; Van Poorten and Post 2005). It is, therefore, possible that the high statewide catch inequality documented in this study is a byproduct of gradual changes in California steelhead population sizes extending beyond the timeframe of the study.

However: abundance is not proportional to CPUE (Maunder et al. 2006; Dedual and Rohan 2016), in part due to ubiquitous phenomena such as catch inequality. These relationships often exhibit hyperstability (Hilborn and Walters 1992), especially when skill levels are high and/or advanced technology is involved (Ward et al. 2013(a); Feiner et al. 2020). Hyperstability, if wholly ignored, can lead to flat trends in CPUE over time and possible ‘business-as-usual’ management approaches, even though true population abundance might be declining, sometimes strongly (Dassow et al. 2020; Simonson et al. 2022; Mrnak et al. 2024). Therefore, understanding the presence and strength of hyperstability in systems with high catch inequality should be a priority research question for fisheries managers going forward, particularly when considering utilization of angler data—such as the SRRC —for population estimations and trends.

Here, we develop a three-pronged framework, including (1) power analysis to determine baseline *n* thresholds, (2) detrending interannual Gini coefficients to account for within-threshold sample size changes over time, (3) assessing whether the Gini coefficient is sensitive to response rates. We showed that this framework is useful because the underlying assumptions regarding acquisition of a measured resource differs between economic contexts (where the resource is income) and fisheries contexts (where the resource is fish catch). Consideration of this nuance is important to calculations of catch inequality, and the subsequent applications for managers. Additionally, we provide our calculated thresholds relative to varying sample sizes as a reference point for future studies (Table 2). At a minimum, our list of sample size breakpoints provides one useful benchmark for the kinds of sample sizes that meet minimum standards for estimating Gini. Given that our across-basin minimum sample size was 77, Gini estimates with substantially lower numbers of angler report cards should be scrutinized.

### Conclusions

Studies of catch inequality shed light on the underappreciated, yet crucial angler dynamics of recreational fisheries (Cook et al. 2001; Seekell et al. 2011; Michaeletz et al. 2008; Dedual & Rohan 2016). The goal of quantifying the Gini coefficient and its drivers is not the pursuit of maximally equal catch distributions; such management endeavors are likely to prove fruitless. Instead, catch inequality can be utilized to understand heterogeneity in the distribution of anglers and their behaviors, as well as its subsequent considerations for data interpretation (Johnston et al. 2010; Matsumura et al. 2019; Hunt et al. 2023). For example, traditional regulations often aim to limit catch and harvest (Seekell et al. 2013), but in fisheries demonstrating extreme catch inequality, such regulations affect only the small subset of anglers who actually catch fish. Should high-catch anglers simply fish harder—by spending more days or hours on the water—traditional regulations become ineffective, as total catch or harvest would remain unchanged (Camp et al. 2023); indeed, total catch could even increase. Furthermore, daily limits are often utilized as a point of reference for angler satisfaction (Cook et al. 2001). When not reached, as is often the case in systems with LACS <1, angler satisfaction may be lower (Cook et al. 2001) and work against angler recruitment or fisheries management efforts. A final yet crucial consideration illustrated by catch inequality is that of hyperstability, especially in highly unequal systems with congregating species such as *O. mykiss* (Dedual and Rohan 2016; Feiner et al. 2020). In these cases, high Gini coefficients may be useful as an indicator of the presence of hyperstability. Therefore, biologists and managers should exercise caution when using CPUE as a proxy for species abundance in these fisheries, as extreme catch inequality can mask overall population declines (Erisman et al. 2011; Dedual and Rohan 2016; Feiner et al. 2020).

## Acknowledgements

We thank the many CDFW biologists and coordinators who have contributed to the collection, preparation, and distribution of the Steelhead Report and Restoration Card Program dataset utilized in this study. We thank the following individuals for their dataset assistance: Erin Fergunson, Daniel Martinez, Aaron Neely, and Jonathan Nelson. We additionally thank Nicholas Gallo for statistical support and Colby Hause for further data curation support. We are grateful to The Fly Fishers of Davis for their support of this research. Finally, we thank the anonymous reviewers for their valuable feedback on our manuscript.

## Competing Interests

The authors declare there are no competing interests.

## Author Contribution Statements

Conceptualization: SRS, CS, ALR

Data curation: SRS

Formal analysis: SRS, ALR, CS

Funding acquisition: SRS, ALR, NAF, RL

Investigation: SRS, ALR

Methodology: SRS, ALR, CS

Project administration: SRS, ALR, NAF, RL

Resources: ALR

Software: SRS

Supervision: ALR, NAF, RL

Validation: SRS, ALR, CS

Visualization: SRS, ALR

Writing – original draft: SRS, ALR

Writing – review & editing: SRS, CS, ALR, NAF, RL

## Funding

S.R.S. was supported by the Fly Fishers of Davis. This research was also supported by the University of California, Davis Agricultural Experiment Station [Grant Nos. CA-D-WFB-2098-H (N.A.F.) and CA-D-WFB-2467-H (A.L.R.)], and the California Trout and Peter B. Moyle Endowment for Coldwater Fish Conservation (A.L.R.).

## Data Availability

The data and code underlying this article are available in Zenodo at DOI <u>10.5281/zenodo.17874424</u>. Only the utilized subset of the larger 2012-2022 CDFW Steelhead Report and Restoration Card Program (SRRC) dataset provided. Full SRRC datasets available upon request from the CDFW SRRC Program.

## Notes

https://doi.org/10.5281/zenodo.17874472

### Competing Interest Statement

The authors have declared no competing interest.

### Summary of Updates

Revisions following reviewer feedback and preparation for publication via CJFAS.

