## Supplemental Materials for "Quantifying catch inequality in recreational fisheries: a case study with California steelhead (*Oncorhynchus mykiss*)"

**Supplementary Materials**

Figure S1. Example of the current Steelhead Report and Restoration Card (SRRC). Provided by the California Department of Fish and Wildlife.

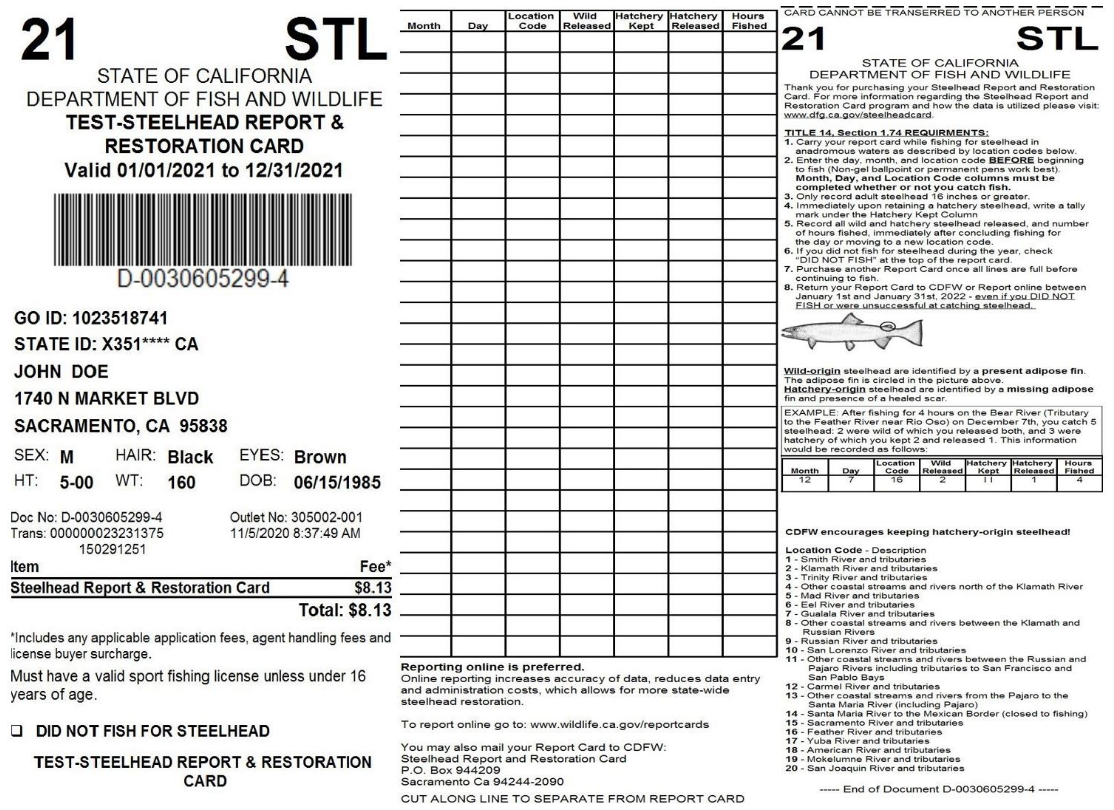

| Table S1. List of United States Geological Survey flow gauges used for flow analysis. | | | | |
| --- | --- | --- | --- | --- |
| **Basin** | **USGS gauge ID** | **Latitude (**$\boldsymbol{}^{\boldsymbol{\circ}}\mathbf{N)}$ | **Longitude (**$\boldsymbol{}^{\boldsymbol{\circ}}\mathbf{W)}$ | **Data range** |
| Smith River | 11532500 | 41.8372657 | 124.0338483 | 1932 - 2024 |
| Klamath River | 11523000 | 41.7054555 | 122.9475025 | 1928 - 2024 |
| Trinity River | 11527000 | 40.9771235 | 123.1415502 | 1932 - 2024 |
| Mad River | 11480390 | 40.4368025 | 123.4812951 | 1980 - 2024 |
| Eel River | 11473900 | 40.575483 | 124.228549 | 1966 - 2024 |
| Coastal R. between Kla. R & Rus. R. ᵃ | 11468000 | 39.170556 | 123.666944 | 1951 - 2024 |
|  | 11468500 | 39.428333 | 123.736667 | 1950 - 2024 |
|  | 11469000 | 40.313333 | 124.282222 | 1912 - 2023 |
| Russian River | 11467000 | 38.711161 | 122.95662 | 1940 - 2024 |
| San Lorenzo River | 11161000 | 37.096363 | 122.072352 | 1953 - 2024 |
| Sacramento River | 11425500 | 39.662492 | 122.024838 | 1946 - 2023 |
| Yuba River | 11421000 | 39.164447 | 121.553488 | 1944 - 2024 |
| American River | 11446500 | 38.637377 | 121.331145 | 1905 - 2024 |
| ᵃ Data averaged across three gauges given cross-basin nature of the basin code. | | | | |

Table S2. Gini coefficients, ranges, and Lorenz Asymmetry Coefficients across all study basins and years, rank-sorted by latitude. Statewide average represents all study basins, 2012-2022. Mean % zero-catch anglers represents annual average percent of anglers catching zero fish at given basin; for Statewide average, represents annual average percent of anglers catching zero fish in all basins.

| **Region** | **Basin** | **Mean**  **Gini** | **Gini**  **range** | **Mean**  **LAC** | **Mean % zero-catch anglers** | **n** |
| --- | --- | --- | --- | --- | --- | --- |
| Statewide | *Average* | *0.81* | *0.69 - 0.91* | *0.88* | 45.8% | *115926* |
| North Coast | *Average* | *0.80* | *0.69 - 0.91* | *0.88* | 52.8% | *76095* |
|  | Smith R. | 0.83 | 0.80 - 0.88 | 0.91 | 57.4% | 8202 |
|  | Klamath R. | 0.75 | 0.71 - 0.80 | 0.90 | 37.0% | 17390 |
|  | Trinity R. | 0.74 | 0.69 - 0.76 | 0.88 | 40.8% | 22520 |
|  | Mad R. | 0.83 | 0.79 - 0.87 | 0.89 | 56.2% | 7155 |
|  | Eel R. | 0.79 | 0.76 - 0.83 | 0.83 | 56.4% | 7268 |
|  | Coastal R. between Kla. R & Rus. R. | 0.80 | 0.78 - 0.83 | 0.91 | 53.3% | 4897 |
|  | Russian R. | 0.85 | 0.81 - 0.91 | 0.84 | 68.7% | 8663 |
| Central Valley | *Average* | *0.83* | *0.75 - 0.90* | *0.91* | 57.0% | *38300* |
|  | Sacramento R. | 0.84 | 0.79 - 0.89 | 0.91 | 58.4% | 13768 |
|  | Feather R. | 0.79 | 0.75 - 0.86 | 0.90 | 46.8% | 7569 |
|  | Yuba R. | 0.85 | 0.81 - 0.87 | 0.90 | 60.1% | 4356 |
|  | American R. | 0.86 | 0.83 - 0.90 | 0.91 | 64.2% | 10838 |
|  | Mokelumne R. | 0.82 | 0.75 - 0.90 | 0.93 | 55.4% | 1769 |
| Other | San Lorenzo R. | 0.82 | 0.76 - 0.89 | 0.79 | 66.7% | 1531 |
